# Programming of neural progenitors of the adult subependymal zone towards a glutamatergic identity by *Neurogenin2*

**DOI:** 10.1101/171686

**Authors:** Sophie Péron, Leo M Miyakoshi, Monika S Brill, Felipe Ortega, Marisa Karow, Sergio Gascón, Benedikt Berninger

## Abstract

While the adult subependymal zone (SEZ) harbors pools of distinct neural stem cells that generate different types of GABAergic interneurons, a small progenitor population in the dorsal SEZ expresses *Neurog2* and gives rise to glutamatergic neurons. Here we investigated whether SEZ progenitors can be programmed towards glutamatergic neurogenesis through forced expression of *Neurog2.* Retrovirus-mediated expression of *Neurog2* induced the glutamatergic neuron lineage markers Tbr2 and Tbr1 in cultured SEZ progenitors which subsequently differentiated into functional glutamatergic neurons. Likewise, retrovirus-mediated expression of *Neurog2* in dividing SEZ progenitors within the adult SEZ induced Tbr2 and Tbr1 expression, hallmarking entry into the glutamatergic lineage also *in vivo.* Intriguingly, *Neurog2*-expressing progenitors failed to enter the rostral migratory stream (RMS) and instead differentiated directly within the SEZ or the adjacent striatum. In sharp contrast, lentivirus-mediated postmitotic expression of *Neurog2* failed to reprogram early SEZ neurons, which instead maintained their GABAergic identity and migrated along the RMS towards the olfactory bulb. Thus, our data show that *Neurog2* can program SEZ progenitors towards a glutamatergic identity, but fails to reprogram their postmitotic progeny.

**Summary statement:** Our study identifies a critical developmental time window during which progenitors of the adult subependymal zone, specified for generating GABAergic neurons, can be reprogrammed towards glutamatergic neurogenesis.

## INTRODUCTION

Accruing evidence indicates that neural stem cells (NSCs) lining the walls of the lateral ventricle in the postnatal and adult subependymal zone (SEZ) exhibit regional identity thereby conferring specific fate restrictions on NSCs (Azim et al., 2015; Lim and Alvarez-Buylla, 2014). Due to this characteristic mosaic organization of the SEZ, NSCs residing in different SEZ domains along the rostro-caudal and dorsal-ventral axes generate neurons of distinct subtype identities and become subsequently destined for distinct sub-domains within the olfactory bulb (OB) (Brill et al., 2009; Merkle et al., 2014; Merkle et al., 2007; Sequerra, 2014). Grafting experiments indicate that these region-specific identities do not become erased upon placing NSCs into heterotopic locations, arguing that extrinsic signals provided locally are not sufficient to reprogram the fate restrictions of NSCs (Merkle et al., 2007). Furthermore, regional fate restrictions appear to extend even to the decision between neuronal and glial fates as indicated by the fact that oligodendrogliogenic NSCs are enriched in the dorsal SEZ and and *in vitro* clearly constitute a lineage distinct from neurogenic NSCs (Ortega et al., 2013).

While the majority of NSCs from the adult SEZ give rise to several types of GABAergic or tyrosine hydroxylase-expressing interneurons (Lim and Alvarez-Buylla, 2014), previous work has shown that a small subpopulation of NSCs located in the dorsal SEZ can generate juxtaglomerular glutamatergic neurons (Brill et al., 2009). This subpopulation is characterized by sequential expression of Pax6, Neurog2, Tbr2, and Tbr1 (Brill et al., 2009) that characterizes glutamatergic neuron producing lineages throughout the forebrain (Hevner et al., 2006). Forced transcription factor expression can alter fate restrictions of neural cells beyond the stem cell stage (Arlotta and Berninger, 2014). Forced expression of *Pax6* (Hack et al., 2005), *Dlx2* (Brill et al., 2008), and *Fezf2* (Zuccotti et al., 2014) has been shown to shift subtype specification within the adult SEZ *in vivo.* When cultured under neurosphere-conditions (high concentration of epidermal growth factor (EGF) and fibroblast growth factor-2, (FGF2)), adult SEZ stem and progenitor cells could be directed towards generation of fully functional glutamatergic neurons by retrovirus-mediated expression of *Neurog2* (Berninger et al., 2007b). Moreover, upon transplantation into the adult hippocampal dentate gyrus, *Neurog2*-expressing adult SEZ stem or progenitor cells exhibited morphological similarities to endogenous dentate granule neurons and some degree of functional integration (Chen et al., 2012). However, exposure to EGF and FGF2 exerts dramatic effects on NSCs that may include a partial loss of regional specification (Gabay et al., 2003; Hack et al., 2004) and may render these cells more permissive to *Neurog2.* Thus, in the present study we addressed the question whether *Neurog2* can overcome fate restrictions of adult SEZ stem and progenitor cells in the absence of elevated growth factor signaling and drive these towards acquisition of a glutamatergic phenotype, and if so, whether such effect extends into the postmitotic life of adult-generated SEZ-derived neurons.

## RESULTS

### Forced expression of *Neurog2* programs adult SEZ progenitors towards a glutamatergic identity *in vitro*

Given that adult-generated glutamatergic OB neurons originate from progenitors located in the dorsal part of the SEZ that express the proneural gene *Neurog2* (Brill et al., 2009) we wondered whether ectopic expression of this transcription factor can program progenitors from the ventral SEZ, which otherwise give rise exclusively to GABAergic neurons, towards a glutamatergic neuron identity. To address this question we first took advantage of an adherent culture of the adult SEZ (Costa et al., 2011; Ortega et al., 2011), which preserves the ratio of GABAergic and glutamatergic neurogenesis as observed *in vivo* (Brill et al., 2009) and transduced proliferating progenitors with a retrovirus encoding *Neurog2*, followed by the reporter *DsRed* behind an internal ribosomal entry site (IRES; Fig. 1A). While cultures infected with a control virus contained only very few neurons (identified by MAP2) expressing the vesicular glutamate transporter-1 (vGluT1), forced expression of *Neurog2* resulted in a massive up-regulation of vGluT1 (20-30 days post infection, DPI), localized to puncta presumably reflecting presynaptic terminals (Fig. 1B). To confirm an actual glutamatergic re-specification, we directly assessed whether *Neurog2*-expressing neurons exhibit glutamatergic synaptic transmission. To this end, we performed pair recordings of neurons derived from *Neurog2*-expressing progenitors and non-transduced control neurons 4 weeks after transduction (Fig. 1C). While synaptic transmission mediated by control neurons was GABAergic in nature (Fig. 1C) as described previously (Costa et al., 2011), stimulation of *Neurog2*-expressing cells in these pairs resulted in postsynaptic currents that were fully blocked by the 2-amino-3-(3-hydroxy-5-methyl-isoxazol-4-yl)propanoic acid (AMPA)/kainate receptor antagonist 6-cyano-7-nitroquinoxaline-2,3-dione (CNQX), demonstrating its glutamatergic nature (Fig. 1C) (n= 6 pairs analyzed). This data demonstrates that forced expression of *Neurog2* can program adult SEZ progenitors towards a glutamatergic identity.

**Figure 1.**
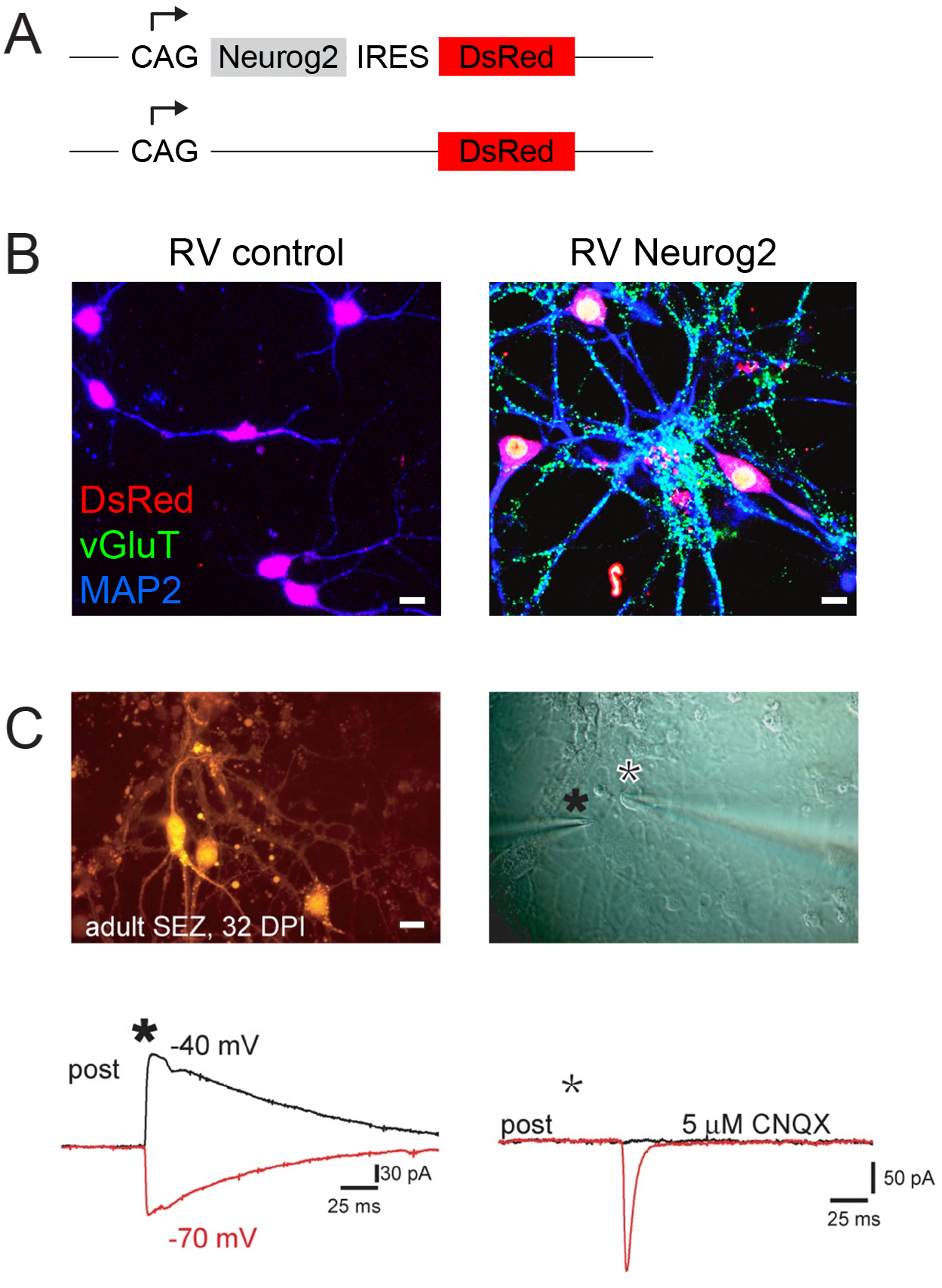
Retrovirus-mediated expression of *Neurog2* induces a glutamatergic phenotype in adult SEZ progenitors. **(A)**, Scheme of the retroviral vectors (RV) CAG-*Neurog2*-lRES-*DsRed* and control CAG-lRES-*DsRed* used in this study. **(B)**, Immunocytochemistry of SEZ cultures 30 days after transduction with the RV CAG-*Neurog2*-lRES-*DsRed* or the corresponding control RV CAG-lRES-*DsRed*. Note that *Neurog2*-expressing cells (right panel; red) are immunopositive for vGluT (green) and MAP2 (blue), indicating a neuronal glutamatergic phenotype. In contrast, cells infected with the *DsRed*-only-encoding vector (left panel; red) remain negative for vGluT. **(C)**, Patch-clamp recording revealing the glutamatergic identity of neurons derived from *Neurog2*-expressing SEZ progenitors. Micrographs show *Neurog2*-expressing cells as indicated by the DsRed fluorescence (left panel) and the recording configuration (right panel). The cell marked with the black asterisk expresses *Neurog2* (DsRed-positive in the left panel), while the cell marked by the white asterisk is untransduced (DsRed-negative in the left panel). Following step depolarisation of the untransduced cell a postsynaptic response is elicited in the *Neurog2*-expressing cell that reversed at a membrane potential more negative than −40 mV characteristic of GABAergic transmission and consistent with the GABAergic nature of the untransduced neuron (left trace). In contrast, stimulation of the *Neurog2*-expressing neuron resulted in a postsynaptic response that was abolished by the AMPA/kainate receptor antagonist CNQX, demonstrating the glutamatergic neuron identity of the *Neurog2*-expressing neuron (right trace). Scale bars: 10 μm (B), 20 μm (C).

### *Neurog2*-mediated fate conversion of adult neural stem cell progeny *in vivo*

Next, we investigated the effects of forced *Neurog2* expression in the progeny of adult SEZ NSCs *in vivo.* To this end, we stereotactically injected retroviruses encoding *DsRed* only, for control, or *Neurog2*-IRES-*DsRed* as experimental manipulation into the adult SEZ. As expected, after seven days, control virus infection revealed the characteristic picture of retrovirally labeled cells migrating in chains along the entire extent of the rostral migratory stream (RMS) towards the core of the olfactory bulb (OB) (Lois et al., 1996). Indeed, 44 ± 7% of cells were found in the SVZ, 19 ± 10% of cells were migrating in the RMS and 37 ± 9% of cells reached the OB (n= 3 mice, 10197 cells analyzed, Fig. 2A-D and I). The DsRed-positive cells in the SEZ (Fig. 2B) and the RMS (Fig. 2C) exhibited morphologies of migratory neuroblasts and upon reaching the OB dispersed radially to establish themselves within the granular and glomerular layer (Fig. 2D). In sharp contrast, progenitors expressing *Neurog2* exhibited a drastically altered migratory behavior (n= 3 mice, 823 cells analyzed), abandoning chain migration (1.5 ± 1% of cells in the RMS) (Fig. 2E,G,I). As a consequence, only extremely few cells finally reached the OB (2 ± 2%) (Fig. 2E,H,I), while most of them remained in the SEZ area (96 ± 1%) (Fig. 2E,F,I). Moreover, although the overall proportion of DCX-positive cells did not differ between control and *Neurog2*-expressing cells (Fig. 3A,B and E-G), *Neurog2*-expressing neurons differentiated within the SEZ or the adjacent striatum, with a highly complex dendritic arborization and high density of dendritic spines (Fig. 3C,D) characteristic of projection neurons. None of these morphologies were observed upon injection of the control virus. To further assess whether *Neurog2*-transduced cells acquire a molecular glutamatergic neuron identity *in vivo*, we stained for the T-box transcription factors Tbr2 and Tbr1, hallmarks of glutamatergic neurogenesis (Hevner et al., 2006). Remarkably, Tbr2 was found to be expressed in 36 ± 8% of *Neurog2*–expressing neurons (n= 3 mice, 7 DPI, 95 cells analyzed) located in the ventral SEZ, which normally is devoid of cells expressing this transcription factor (0.7 ± 0.3% of Tbr2-positive cells among cells infected with the control retrovirus, n= 3 mice, 7 DPI, 962 cells analyzed) (Fig. 3H-J). The acquisition of a glutamatergic neuron program by *Neurog2*-expressing cells was further corroborated by the presence of a similar proportion of Tbr1-positive cells (n= 2, 7 DPI, 215 cells analyzed) (Fig. 3K).

**Figure 2.**
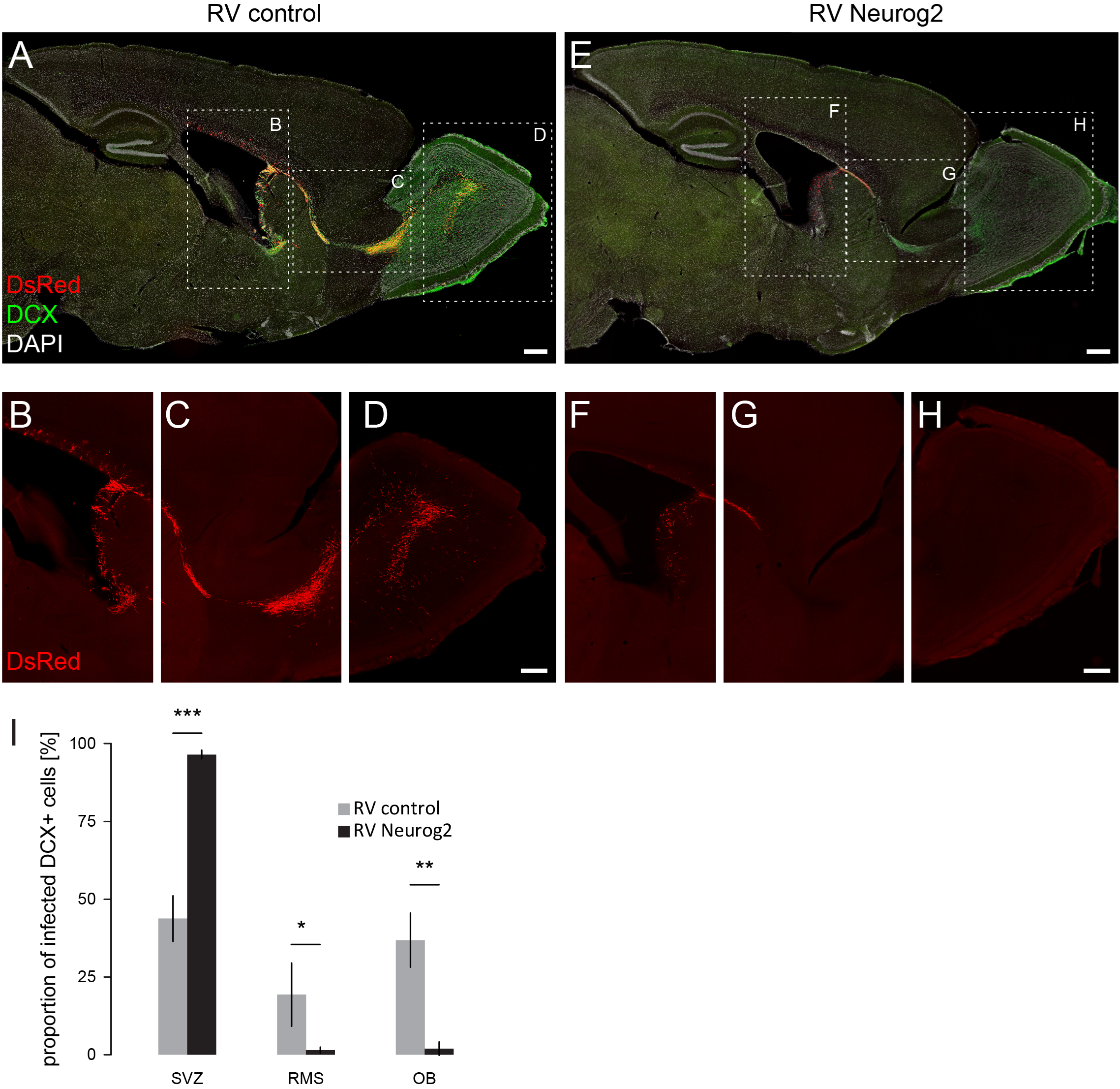
Retrovirus-mediated *Neurog2* expression *in vivo* alters the migration behaviour of SEZ progenitors. **(A-D)**, Sagittal view of an adult mouse brain depicting SEZ cells transduced with the control RV CAG-lRES-*DsRed* (red) (B), migrating through the RMS (C) and reaching the OB (D). **(E-H)**, Micrographs showing cells transduced with the RV CAG-*Neurog2*-lRES-*DsRed* (red). At 7 DPl, only few transduced neuroblasts partially entered the RMS (G) and failed to reach the core of the OB (H). They instead remained stationary in the anterior portion of the SEZ (F). **(I)**, Quantification of the number of DCX-positive/DsRed-positive infected cells in the SVZ, RMS and OB at 7 DPl. Error bars indicate mean ± s.d., n=3/group. ^∗^p^<^0.05, ^∗∗^p^<^0.01, ^∗∗∗^p^<^0.001, One-Way ANOVA followed by Tukey’s HSD post-hoc test. Scale bars: 1 mm (A,E), 500 μm (B-D and F-H).

**Figure 3.**
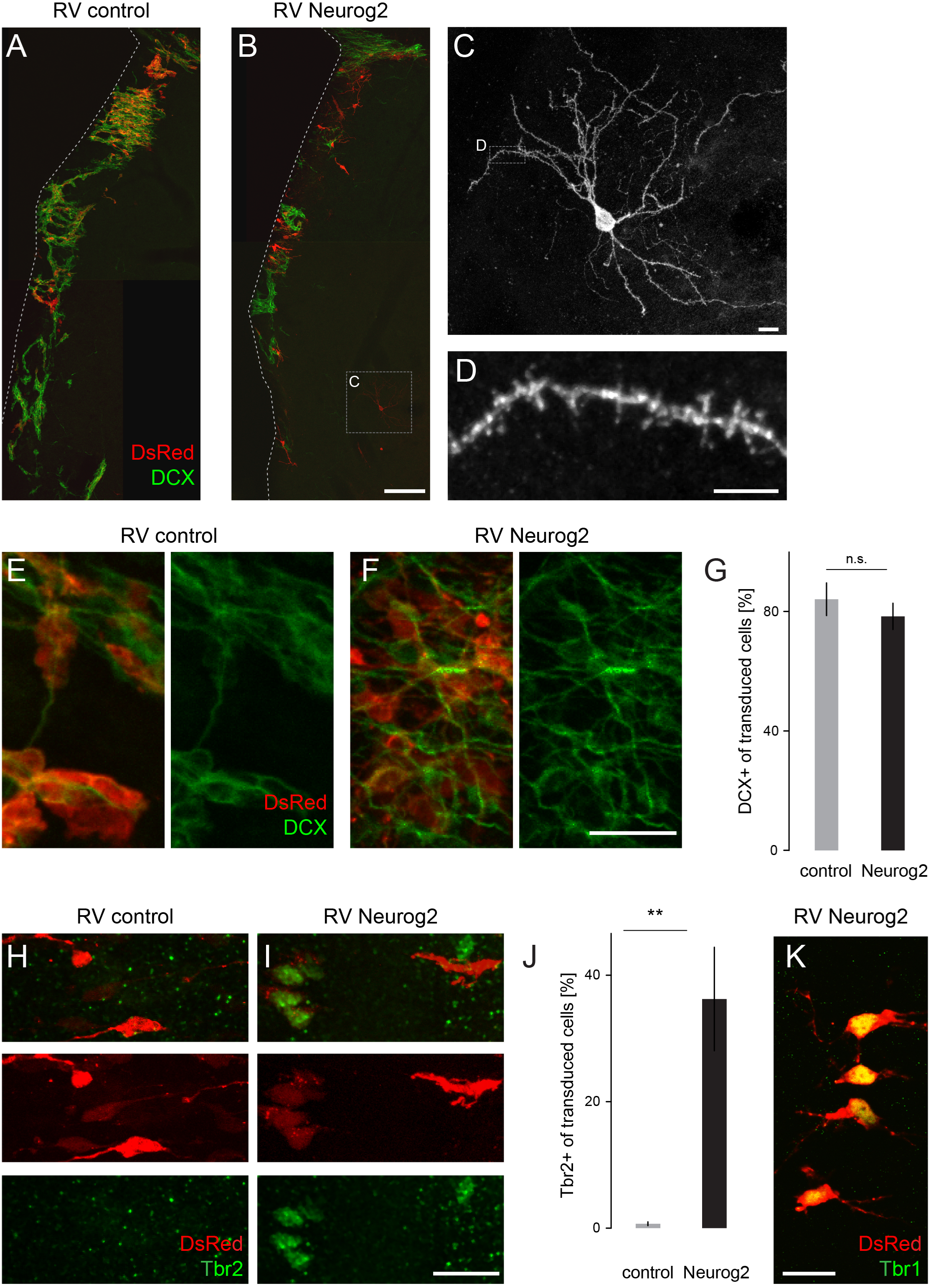
Retrovirus-mediated *Neurog2* expression induces a glutamatergic neuron identity *in vivo.* **(A,B)**, Micrographs depicting transduced SEZ cells with either control RV (red) (A) or RV CAG-*Neurog2*-lRES-*DsRed* (red) (B) co-expressing DCX (green) at 7 DPl. Note that some *Neurog2*-transduced neuroblasts invade the adjacent striatum. **(C,D)** Higher-power images showing a *Neurog2*-transduced cell with neuronal morphology (C) and spines (D). **(E,F),** Micrographs depicting transduced cells immune-positive for DCX (green) **(G),** Quantification of the percentage of DCX-positive cells among infected cells. Error bars indicate mean ± s.d., n.s, non-significant, t-test. **(H-I),** Micrographs showing the expression of the transcription factor Tbr2 (green) in SEZ cells forced to express *Neurog2* (I), while cells infected with the control retrovirus lack Tbr2 expression (H). **(J),** Quantification shows that more than one-third of *Neurog2*-expressing cells up-regulated Tbr2. Error bars indicate mean ± s.d., n=3/group. ^∗∗<^0.01, t-test. **(K),** Micrograph depicting Tbr1 expression (green) in cells transduced with the RV CAG-*Neurog2*-IRES-*DsRed* (red). Scale bars: 100 μm (A,B), 10 μm (C), 5 μm (D), 25 μm (E,F), 20μm (H,I,K).

### Forced expression of *Neurog2* in young postmitotic neurons does not result in reprogramming towards a glutamatergic phenotype

In the above experiments, use of retroviral vectors encoding *Neurog2* restricted transduction to fast-dividing cells, most of which are transit-amplifying precursors (Costa et al., 2011; Doetsch et al., 2002). We next asked whether conversion towards a glutamatergic identity would be possible at later stages of lineage progression, i.e., after the last cell division when these cells become postmitotic and commence differentiation.

Thus, we employed lentiviral vectors encoding *Neurog2* and *egfp* allowing for transduction of non-dividing cells. In these constructs, expression of *Neurog2* and the reporter are driven by the minimal human synapsin promoter (hSyn, Fig. 4A) restricting expression of *Neurog2* and *egfp* to postmitotic neurons (Gascon et al., 2008). In fact, transduction of primary cortical cultures from embryonic day E14 with the lentiviral vector hSyn-*Neurog2*-IRES-*egfp* resulted in efficient expression of Neurog2 and GFP proteins, which were both restricted to Tuj-1-immunoreactive neurons (Fig. 4B). We then examined whether hSyn-driven expression of *Neurog2* in adult SEZ cultures causes a similar conversion towards a glutamatergic identity as observed following transduction at the precursor stage. In sharp contrast to the effect of *Neurog2* observed in dividing progenitors, the vast majority of the lentivirus-transduced neurons remained immunostaining-negative for vGluT1 at 30 days post infection (Fig. 4C), although vGluT1 positive neurons were readily detectable (Fig. 4D). However, their number did not differ from cultures transduced with a control lentiviral vector (2.5% vs 4%, 701 cells analyzed) (Fig. 4E), indicating that these few glutamatergic neurons represent the small endogenous glutamatergic population derived from the dorsal portion of the adult SEZ (Brill et al., 2009). This data show that forced *Neurog2* expression fails to convert immature postmitotic neurons derived from adult NSCs into glutamatergic neurons *in vitro.*

**Figure 4.**
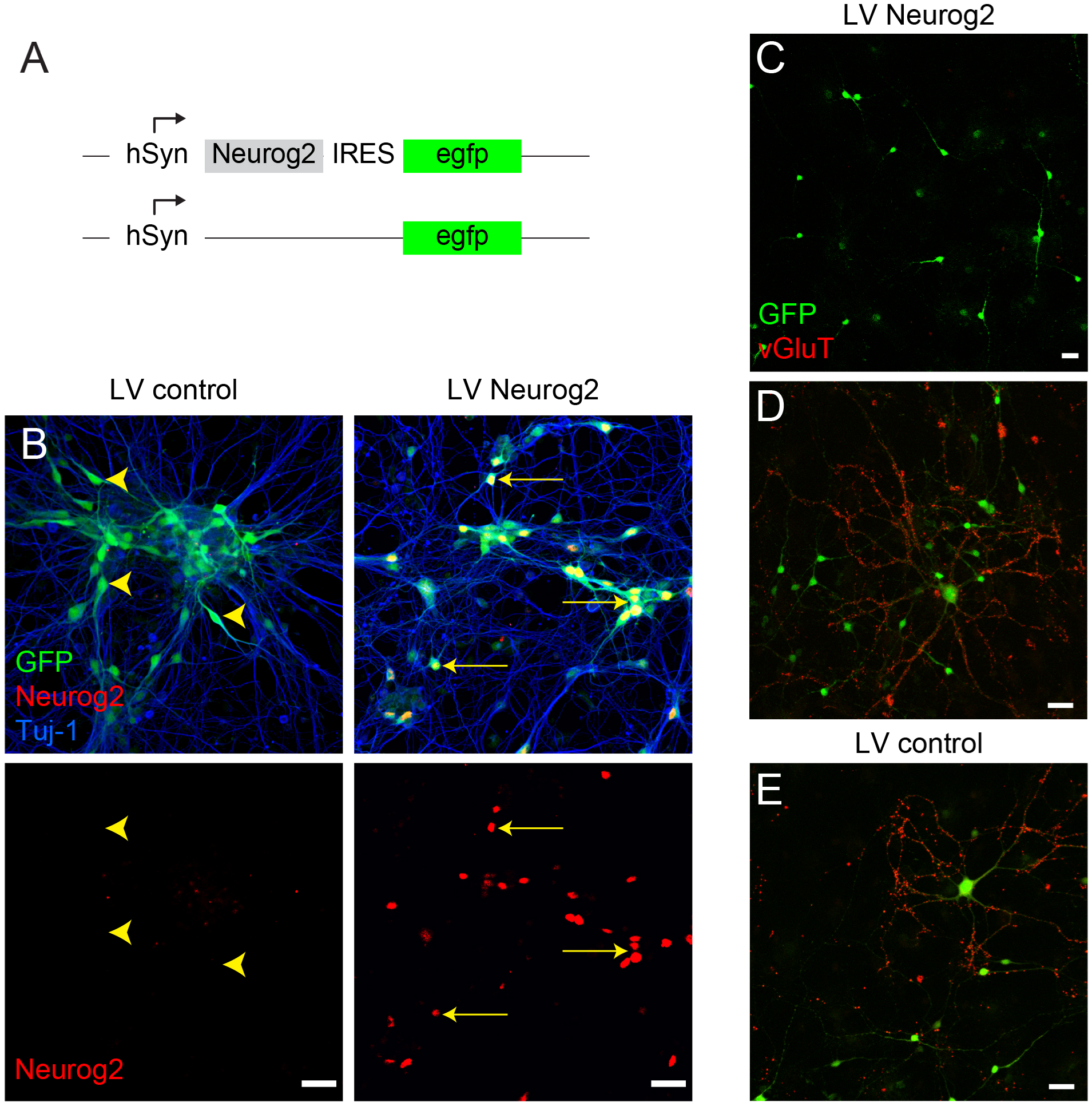
Forced *Neurog2* expression fails to induce a glutamatergic phenotype in postmitotic neurons derived from SEZ progenitors. **(A)**, Scheme of lentiviral vectors hSyn-*Neurog2*-IRES-*egfp* and hSyn-*egfp* used. **(B)**, Micrographs show the expression of Neurog2 protein (red) and GFP (green) in primary cortical cultures from embryonic day E 14, 6 days after transduction with the hSyn-*Neurog2*-IRES-*egfp* lentiviral vector (right panel), or the control lentiviral vector encoding only GFP (left panel). Note that the human synapsin (hSyn) promoter drives selective expression of the transgene in Tuj-1 neurons (blue). **(C)**, In adult SEZ primary cultures, most of the neuroblasts transduced with the Syn-*Neurog2*-IRES-*gfP* do not differentiate into vGluT-positive (red) glutamatergic neurons at 30 DPI. **(D,E)** Example micrographs of *Neurog2*-expressing neurons (hSyn-*Neurog2*-IRES-*egfP*) (D) and control neurons (hSyn-*egfP*) (E) exhibiting vGluT immunoreactivity, indicative of the rare presence of endogenous glutamatergic neurons. Scale bars: 50 μm (B-E).

### *Neurog2* activates the transcriptional program of glutamatergic neurogenesis in SEZ progenitors, but fails to do so in young postmitotic neurons

Forebrain glutamatergic neurogenesis is generally characterized by the sequential expression of Tbr2 and Tbr1 (Hevner et al., 2006) and previous work has delineated the same expression sequence in glutamatergic neurons derived from *Neurog2*-expressing SEZ progenitors (Brill et al., 2009). Thus, we examined whether forced *Neurog2* expression results in the activation of the same molecular pathway in adult SEZ progenitor cells. To this end, we transduced cultured SEZ progenitors with the retrovirus encoding *Neurog2* and analyzed Tbr2 and Tbr1 expression at days 7 and 9 post infection, respectively. Consistent with the natural program of glutamatergic neurogenesis we observed expression of both transcription factors in *Neurog2*-expressing cells (Fig. 5A,B). Tbr2 was expressed only in a subpopulation of *Neurog2*-transduced cells (44 ± 3%; Fig. 5D), most likely due to the fact that this transcription factor is expressed transiently during glutamatergic lineage progression (Hevner et al., 2006). We next transduced adult SEZ cultures with the LV-hSyn-*Neurog2*-IRES-*egfp* Contrary to the effect of retroviral expression at the progenitor stage, expression of *Neurog2* in immature postmitotic neurons did not induce Tbr2 (Fig. 5A,D) or Tbr1 (data not shown) expression. This was more conspicuous as Tbr2 was found to be expressed in a minor subpopulation of untransduced SEZ cells (5 ± 5%; Fig. 5A), again reflecting the low degree of intrinsic glutamatergic neurogenesis from adult SEZ progenitors (Brill et al., 2009). Finally, in agreement with the failure to activate a glutamatergic program, neurons transduced with LV-hSyn-*Neurog2*-IRES-*egfp* maintained GABA immunoreactivity (94 ± 6%; Fig. 5C,D), while retrovirus-mediated *Neurog2* expression at the progenitor stage resulted in a loss of GABA immunoreactivity in line with the acquisition of a glutamatergic phenotype (Fig. 5C,D). Thus, this data not only confirm that *Neurog2*-transduced SEZ progenitors undergo a fate conversion, but that this involves the recapitulation of glutamatergic neurogenesis from adult SEZ progenitors under physiological conditions (Brill et al., 2009). On the other hand, the failure of inducing a glutamatergic phenotype in young postmitotic neurons following forced *Neurog2* expression is also accompanied by failure of inducing the stereotypical program of glutamatergic lineage progression.

**Figure 5.**
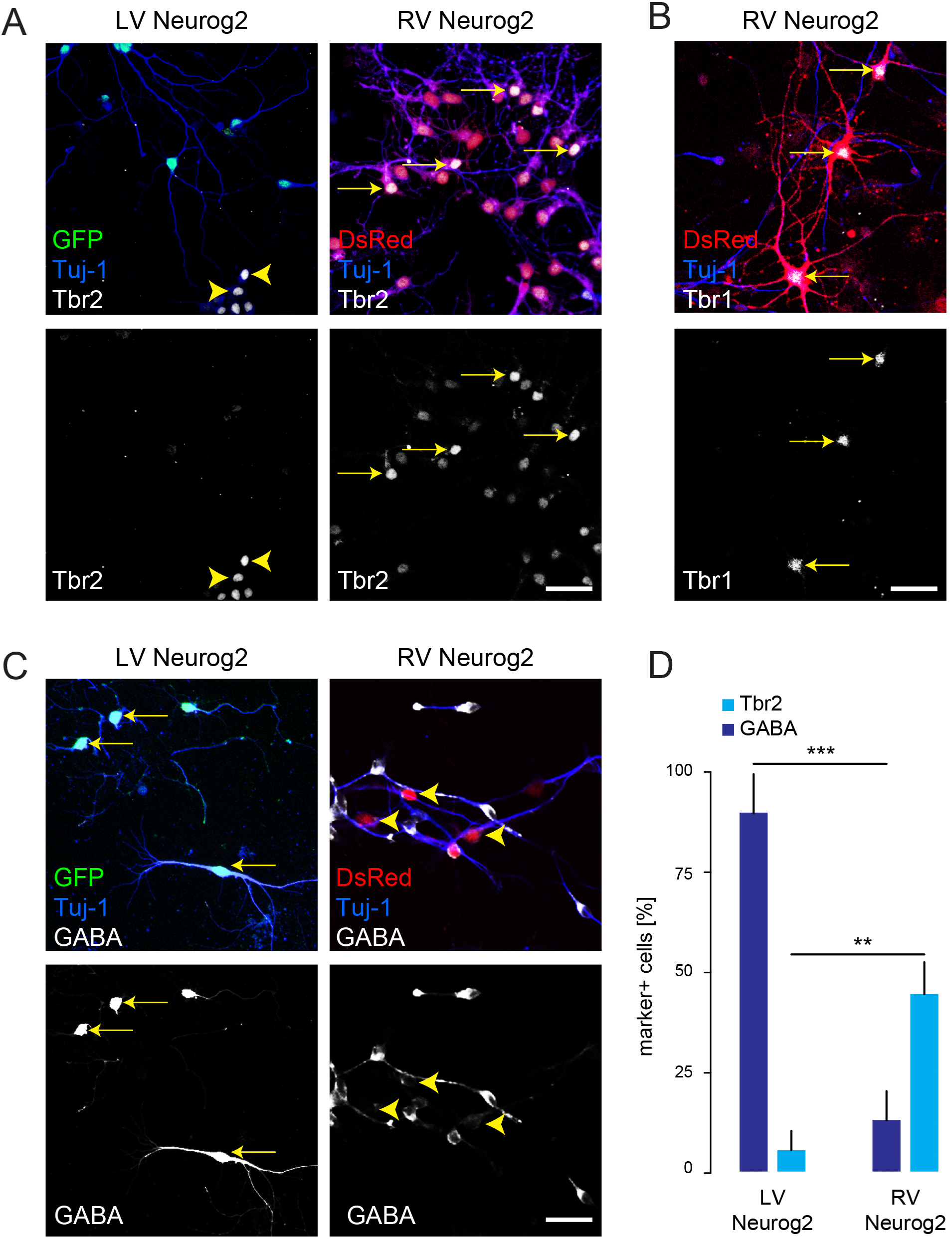
*Neurog2* induces hallmarks of the glutamatergic lineage when expressed in dividing progenitors but not in young postmitotic neurons. **(A)**, Cells from SEZ cultures infected with the RV CAG-*Neurog2*-IRES-*DsRed* differentiate into Tuj-1-positive neurons expressing the early glutamatergic transcription factor Tbr2 (right panel, arrows) at 7 DPI, while those cells infected with the lentiviral hSyn-*egfP* control vector fail to express Tbr2 (left panel). Note the cluster of non-transduced progenitors expressing Tbr2 (arrowheads) occurring at low frequency in SEZ cultures. **(B)**, At later time points (9 DPI), neurons derived from progenitors transduced with RV CAG-*Neurog2*-IRES-*DsRed* also express Tbr1 (arrows). **(C)**, Tuj-1-positive SEZ neuroblasts transduced with RV CAG-*Neurog2*-IRES-*DsRed* are devoid of GABA immunoreactivity at 7 DPI (right panel, arrowheads) while neuroblasts transduced with the lentiviral vector Syn-*neurog2*-IRES-*egfp* remain immunoreactive for GABA (left panel, arrows). **(D)**, Quantification of the proportions of transduced cells immunoreactive for Tbr2 or GABA at 7 DPI following retro- or lentiviral-mediated expression of *Neurog2.* Error bars indicate mean ± s.d. ^∗∗^p^<^0.01, ^∗∗∗^p^<^0.001, t test. Scale bars: 60 μm (A-C).

### *Neurog2* expression in young postmitotic neurons fails to induce a glutamatergic phenotype *in vivo*

We next aimed to assess the effect of forced *Neurog2* expression in young postmitotic neurons following stereotactic injection of LV-hSyn-*Neurog2*-IRES-*egfp* into the adult SEZ *in vivo* (Fig. 6). In contrast to the aberrant migration of cells transduced with a retrovirus encoding *Neurog2* (Fig. 2), hSyn-driven *Neurog2* expression did not result in alterations in the migration program. Indeed, DCX-positive cells were found to migrate normally via the RMS towards the OB (control: 9 ± 4.5% of cells in SEZ, 3 ± 0.6% of cells in RMS and 88 ± 5% of cells in OB, n= 3 mice, 1929 cells analyzed; *Neurog2*: 10 ± 4% of cells in SEZ, 2.5 ± 1% of cells in RMS and 88 ± 5.5% of cells in OB, n= 3 mice, 855 cells analyzed; Fig. 6A). The presence of DCX-positive cells expressing GFP while leaving the RMS indicates the early activity of the hSyn-promoter in SEZ cells transduced with the LV-hSyn-*Neurog2*-IRES-*egfp* (Fig. 6B). Notably, at 10 DPI a large number of transduced cells were already present in the OB and dispersed radially towards the more superficial layers, where the majority acquired the morphology characteristic of GABAergic granule or periglomerular neurons (Fig. 6C,D). Also at that late stage, we were unable to detect expression of the glutamatergic lineage marker Tbr2 in both, granule or periglomerular neurons, amongst the LV-hSyn-*Neurog2*-IRES-*egfp* cells (Fig. 6E-F). This data indicate that *Neurog2* expression at a postmitotic stage fails to reprogram the phenotype of neurons derived from adult NSCs.

**Figure 6.**
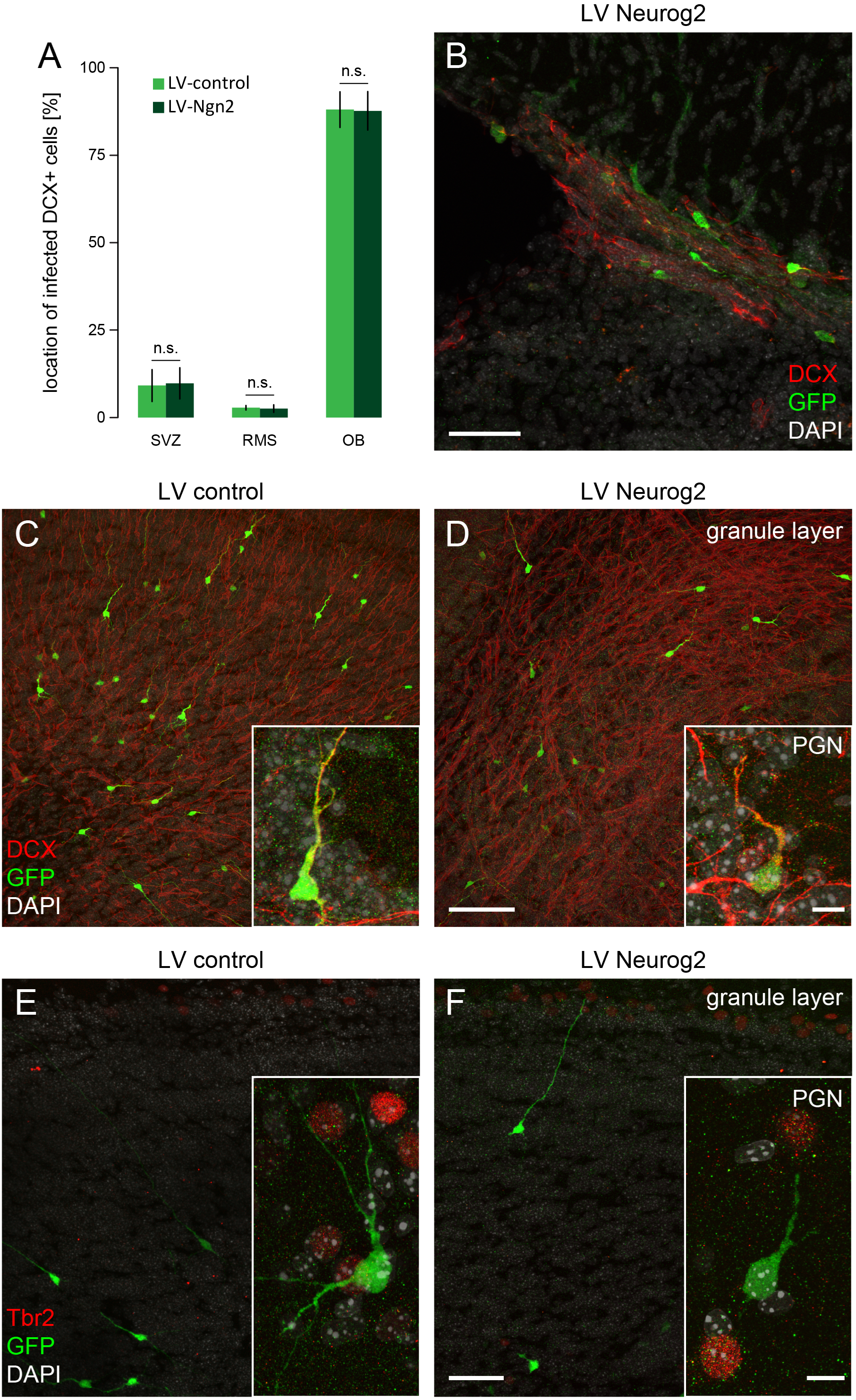
Expression of *Neurog2* fails to induce a glutamatergic phenotype in postmitotic neurons derived from SEZ progenitors *in vivo.* **(A)**, Quantification of the number of DCX-positive/GFP-positive LV-transduced cells in the SVZ, RMS and OB shows that the migration of SEZ cells to the OB is not affected by post-mitotic induction of *Neurog2.* Error bars indicate mean ± s.d., n=3/group, n.s., non-significant, One-Way ANOVA followed by Tukey’s HSD post-hoc test. **(B),** Micrograph showing neuroblasts (DCX, in red) leaving the SEZ and expressing GFP (in green), indicative of lentiviral transduction with *Neurog2* and hSyn-promoter activity in these cells. **(C,D),** Micrographs of the granule layer of the OB showing lentivirally transduced cells (green) migrating radially and integrating as granule and or periglomerular neurons (PGN, insets, DCX red) at 10 DPI. **(E-F),** SEZ cells transduction with the hSyn-*egfp* control (green) and the hSyn-*Neurog2*-IRES-*egfp* (green) lentiviruses do not result in Tbr2 expression (red) in periglomerular (PGN, insets) or granule neurons. Scale bars: 50 μm (B,C,D,E,F), 10 μm (insets in C,D,E,F).

## DISCUSSION

In the present study we demonstrate that forced expression of *Neurog2* redirects the program of proliferating adult SEZ progenitors, mostly giving rise to GABAergic olfactory neurons, towards generating neurons of the glutamatergic lineage. This indicates that *Neurog2* can override region specific fate restrictions of adult NSCs. However, once SEZ-derived cells have differentiated into postmitotic neurons, *Neurog2* can no longer induce a switch in transmitter phenotype, as neurons retain their GABAergic neuron identity. Thus, there appears to be a restricted time window during the lineage progression from stem cell to neuron during which *Neurog2* can alter the program determining transmitter identity. Our data supporting a stage- and hence cellular context-dependent potency of Neurog2 are in line with a recent study showing that gain-of-function of Neurog2 can bias the balance between deep-layer and upper-layer neurogenesis towards the former in early cortical progenitors, but has a limited capacity to specify early neuronal features in late cortical progenitors (Dennis et al., 2017). Our data also provide additional evidence for *Neurog2*’s instructive role for glutamatergic neurogenesis (Mattar et al., 2008).

### Retroviral *Neurog2* expression directs dividing SEZ progenitors towards the glutamatergic neuron lineage

Retroviruses predominantly transduce fast-dividing progenitors both *in vitro* and *in vivo.* Thus, by employing a retroviral vector for delivery of *Neurog2*, expression of the proneural gene should be largely confined to activated stem cells and transit-amplifying progenitors as well as dividing neuroblasts. We found that retrovirus-mediated *Neurog2* expression resulted in the acquisition of a glutamatergic neuron identity *in vitro*, as evidenced by expression of the glutamatergic lineage transcription factors Tbr2 and Tbr1 (Hevner et al., 2006), expression of vesicular glutamate transporter and the development of glutamatergic synapses. In accordance with a fate switch *in vivo*, *Neurog2*-expressing cells no longer migrated to the OB via the RMS, but instead differentiated within the SEZ or the adjacent striatum. Of note, a recent study showed SVZ cells can be redirected from their normal migration route and directed towards other brain regions upon co-transduction with retroviruses encoding *Neurog2* and *Isl1* (Rogelius et al., 2008). Thus, part of the fate switch induced by *Neurog2* is an alteration of the migratory program normally under the influence of *Dlx2* (Brill et al., 2008). Besides this, differentiated neurons derived from *Neurog2*-expressing progenitors developed morphological features reminiscent of pyramidal-like neurons. Most importantly a substantial portion of the *Neurog2*-expressing cells co-expressed Tbr2 and Tbr1, demonstrating entry into the glutamatergic neuron lineage (Hevner et al., 2006).

### *Neurog2* fails to reprogram postmitotic neurons derived from SEZ progenitors towards the glutamatergic neuron lineage

In sharp contrast to the conspicuous effect of retrovirus-mediated *Neurog2* expression, targeting *Neurog2* to postmitotic neurons by employing a lentivirus driving transgene expression from the human synapsin promoter (Gascon et al., 2008) failed to alter the already ongoing program of OB interneuron genesis. Lentivirally *Neurog2*-transduced cells continued to migrate along the RMS to the OB where they took position within the granule cell layer for which most of the adult-generated neurons in the SEZ are destined (Merkle et al., 2014; Merkle et al., 2007). Also, in contrast to *Neurog2* expression in progenitors, postmitotic expression did not cause overt changes in morphology and GFP-expressing cells exhibited morphologies highly reminiscent of OB granule neurons. Moreover, primary cultures of neurons derived from SEZ progenitors were GABA immunoreactive when *Neurog2* was expressed postmitotically. Finally, both *in vitro* and *in vivo*, *Neurog2* failed to induce the expression of glutamatergic lineage transcription factors such as Tbr2. In the absence of any evidence for a fate change, it thus appears that *Neurog2*-expressing SEZ-derived neurons maintain the interneuron identity originally acquired during lineage progression from stem cell to neuron. This data argues for the establishment of powerful epigenetics barriers that impede a Neurog2-instructed switch from a GABA-to-glutamatergic transmitter identity.

### Specific windows of opportunity for *Neurog2*-induced programming and reprogramming

This data suggest that the potency of *Neurog2* sharply declines when neurons become postmitotic and is likely linked to changes in the neuronal epigenome during this transition as strongly indicated by the failure to upregulate Tbr2 which is known to be a direct target of Neurog2 (Kovach et al., 2013; Ochiai et al., 2009). Alternatively, or in addition to epigenetic barriers, the effectiveness of Neurog2 action may be curtailed by signaling mechanisms regulating the phosphorylation state (Quan et al., 2016) or the formation of homo-versus heterodimers (Li et al., 2012).

The failure of reprogramming at later stages of the stem cell to neuron lineage progression is even more conspicuous given the capacity of *Neurog2* to sequentially induce Tbr2 and Tbr1 expression in astrocytes derived from the early postnatal cerebral cortex and reprogram these into fully functional glutamatergic neurons (Berninger et al., 2007a; Heinrich et al., 2010). Yet, also this reprogramming activity of proneural genes such as *Neurog2* or *Ascl1* appears to become more restricted with glial maturation (Masserdotti et al., 2015; Ueki et al., 2015) and glia-to-neuron conversion by *Neurog2* or *Ascl1* alone is very limited in the adult central nervous system *in vivo* (Grande et al., 2013). Remarkably, some of these restrictions can be overcome by allowing for enhanced epigenetic remodeling as recently demonstrated for *Ascl1*-induced reprogramming of Müller glia into bipolar neurons in the lesioned adult mouse retina (Jorstad et al., 2017). Moreover, a recent study identified a critical metabolic checkpoint for *Neurog2*- and *Ascl1*-induced glia-to-neuron reprogramming which could be negotiated by co-expression of *Bcl2* (Gascon et al., 2016). While this check point negotiation appears to work predominantly via interference with reactive oxygen species-induced ferroptosis, it is conceivable that enhanced reprogramming following *Bcl2* co-expression might also be related to the need of producing larger quantities of particular mitochondrial metabolites required for epigenetic remodeling such as shown in other systems (Wong et al., 2017).

The developmental window-specific actions of *Neurog2* on adult SEZ stem and progenitor cells exhibit remarkable parallelism and differences to that of the transcription factor *Fezf2. Fezf2* was originally described as a transcription factor to specify the fate of cortical progenitors towards a corticofugal identity during development (Molyneaux et al., 2005). Subsequent work demonstrated that it can reprogram striatal progenitors towards a corticofugal identity *in vivo* (Rouaux and Arlotta, 2010) thus causing not only a similar transmitter identity switch (from GABAergic to glutamatergic neuron) as described here for *Neurog2*, but also eliciting a specific neuronal subtype conversion (from medium spiny neuron to corticofugal pyramidal neuron). Of note, *Fezf2* reprogramming activity extends into early postmitotic life of a neuron but markedly declines in the course of few days (Rouaux and Arlotta, 2013) arguing for the presence of a critical window of nuclear plasticity that closes with epigenetic changes that occur during neuronal maturation (Amamoto and Arlotta, 2014). More recently, *Fezf2* was found to program NSCs in the postnatal and adult SEZ towards a glutamatergic neuron identity *in vitro* and *in vivo* (Zuccotti et al., 2014). In conspicuous difference to the findings reported here for forced *Neurog2* expression in fast-dividing cells, the *Fezf2*-induced fate switch was restricted specifically to the stem cell stage and failed to convert both transit-amplifying progenitors or dividing neuroblasts. Moreover, in contrast to the redirection of *Neurog2*-expressing neurons, progeny of *Fezf2*-expressing NSCs still migrated to the OB. Thus, *Fezf2* and *Neurog2* appear to possess distinct temporal windows of opportunity.

Taken together, our data provide evidence for remarkable plasticity within the lineage of adult NSCs. It will be interesting to learn whether this plasticity can be harnessed towards translational approaches to recruit adult NSC progeny into diseased brain tissue for repair (Benraiss et al., 2013; Brill et al., 2009; Gage and Temple, 2013; Saghatelyan et al., 2004). To fully exploit this, it will be of crucial importance to identify the mechanisms underlying the closure of windows of opportunity and thereby terminally sealing neuron fate.

## MATERIAL AND METHODS

### Ethical approval

All animal procedures were performed in accordance to the Policies on the Use of Animals and Humans in Neuroscience Research, revised and approved by the Society of Neuroscience and the state of Bavaria under license number 55.2-1-54-2531-144/07.

### Plasmids and DNA constructs

Retroviral and lentiviral transduction of SEZ primary cultures was performed 2 hours after plating the cells on glass coverslips, using VSV-G (vesicular stomatitis virus glycoprotein) pseudotyped viruses. For the transduction with retrovirus we used the retroviral vectors RV-pCAG-*Neurog2*-IRES-*DsRed* and RV-pCAG-IRES-*DsRed* as described previously (Heinrich et al., 2011). To obtain the lentiviral vector Syn-*DsRed*-Syn-*egfp* we re-cloned a fragment containing the cDNA of *Neurog2* and the internal ribosomal entry site from the RV-pCAG-*Neurog2*-IRES-*DsRed* into the lentiviral vector hSyn-*DsRed*-Syn-*egfp* (Gascon et al., 2008). In this case, control experiments were performed with the lentiviral vector hSyn-*egfP* (Gascon et al., 2008).

### SEZ primary culture

Following a previously established protocol by (Costa et al., 2011; Ortega et al., 2011), SEZ cultures were prepared from the lateral wall of the lateral ventricle of young adult (8 - 12 weeks) C57/Bl6 mice (*Mus musculus*) (Costa et al., 2011; Ortega et al., 2011). Briefly, tissue was dissociated in 0.7 mg/ml hyaluronic acid (Sigma-Aldrich) and 1.33 mg/ml trypsin (Sigma-Aldrich) in Hanks’ Balanced Salt Solution (HBSS; Invitrogen) with 2 mM glucose (Sigma-Aldrich) at 37°C for 30 minutes. After this enzymatic treatment, an equal volume of an ice-cold medium consisting of 4% bovine serum albumin (BSA; Sigma-Aldrich) in Earle’s Balanced Salt Solution (EBSS; Invitrogen) buffered with 20 mM HEPES (Invitrogen) was added in order to stop dissociation. Cells were then centrifuged at 200 ***g*** for 5 minutes, re-suspended in ice-cold medium consisting of 0.9 M sucrose (Sigma-Aldrich) in 0.5× HBSS, and centrifuged for 10 minutes at 750 ***g***. The cell pellet was re-suspended in 2 ml ice-cold medium consisting of 4% BSA in EBSS buffered with 2 mM HEPES, and the cell suspension was placed on top of 12 ml of the same medium and centrifuged for 7 minutes at 200 ***g***. The resulting cell pellet was resuspended in DMEM/F12 Glutamax (Invitrogen) supplemented with B27 (Invitrogen), 2 mM glutamine (Sigma), 100 units/ml penicillin (Invitrogen), 100 μg/ml streptomycin (Invitrogen), buffered with 8 mM HEPES. Finally, cells were plated on poly-d-lysine (Sigma) coated coverslips at a density of 200-300 cells per mm^2^ and, after 2 hours to allow for settlement of the cells, cultures were treated with retroviral or lentiviral vectors for transduction.

### Viral vector injections

Stereotactic injections of retrovirus and lentivirus were performed in 2 - 3 months old C57BL/6 male mice (*Mus musculus*). Prior to stereotactic injections, mice were anesthetized using Ketamine (100 mg/kg; CP-Pharma) and Xylazine (5 mg/kg; Rompun; Bayer) and placed into a stereotaxic frame. Approximately, 0.5 μl viral suspension was injected using a pulled-glass capillary at the following coordinates relative to Bregma: 0.7 (antero-posterior), 1.2 (medio-lateral) and 2.1-1.7 (dorso-ventral).

### Immunohistochemistry and immunocytochemistry

Mice were deeply anesthetized using 5% chloralhydrate (wt/vol) diluted in phosphate buffered saline and then perfused transcardially with saline (0.9%), followed by 4% paraformaldehyde (PFA) (wt/vol) for 30 minutes. After this initial fixation, brains were dissected and post-fixed for at least 2 hours in 4% PFA. Sagittal brain sections were prepared at a thickness of 50 μm using a Thermo Scientific vibrating blade microtome.

For immunohistochemistry, sections were blocked for 90 minutes in TBS containing 0.3% Triton and 5% donkey serum. Primary antibodies were diluted in blocking solution and incubated over night at 4°C on the sections. In this study, we used antibodies to Tbr1 (Rabbit, 1:100 Abcam, ab31940), Tbr2 (Rabbit, 1:500 Abcam, ab23345), RFP (Rabbit, 1:500 Rockland, 600401379S), Doublecortin (DCX, Goat, 1:300 Santa Cruz Biotechnology, sc-8066) and GFP (Chicken, 1:1000, Aves labs, GFP-1020) The next day samples were washed with TBS and subsequently incubated at room temperature for 1 hour with species-corresponding secondary antibodies made in donkey and conjugated to Cy3 (1:500, Dianova, 711-165-152, 705-165-147), Alexa Fluor 488 (1:200, Invitrogen, A21206, A11055; Jackson Immunoresearch, 703-545-155) and Alexa Fluor 647 (1:500, Invitrogen, A31573, A21447). Sections were washed again in TBS, counterstained with DAPI and mounted with an acqueous mounting medium.

Cultures were fixed in 4% PFA in PBS for 15 minutes at room temperature and processed for antibody staining as described previously (Ortega et al., 2011).

### Electrophysiology

Perforated patch-clamp recordings were performed as previously described (Heinrich et al., 2011).

### Quantitative and statistical analysis

Images stacks were acquired using a confocal microscope (Olympus FV1000 equipped with x20/0.8 N.A., x40/1.35 and x60/1.42 N.A. oil-immersion objectives). Quantifications of marker-positive cells were performed on single optical sections of an image stack. The percentage of DCX-positive or Tbr2-positive cells was calculated among RFP-positive cells in the SEZ region. For distribution analysis of the cells, RFP-positive/DCX-positive cells were quantified in the SEZ, RMS and OB, and the percentage of cells in each region among total number of RFP-positive/DCX-positive cells was calculated. The number of animals (n) used and the total number of cells is indicated in the text. Data are represented as mean ± s.d. Statistical analysis was performed using SPSS Statistics 22 software. The statistical tests used and p values are indicated in the figure legends.

## Acknowledgements

We are grateful to Dr. Magdalena Götz for support throughout the project.

## Author contributions

S.P., L.M.M and M.S.B designed and performed experiments, interpreted and analyzed results, and wrote the manuscript. F.O. and M.K. designed and performed experiments. S.G. conceptualized, designed and performed experiments, interpreted and analyzed results, and wrote the manuscript. B.B. conceptualized the study and experiments, interpreted results, and wrote the manuscript. All authors discussed the manuscript.

